# The Elastic Network Contact Model applied to RNA: enhanced accuracy for conformational space prediction

**DOI:** 10.1101/198531

**Authors:** Olivier Mailhot, Vincent Frappier, François Major, Rafael Najmanovich

## Abstract

**Motivation:** The use of Normal Mode Analysis (NMA) methods to study both protein and nucleic acid dynamics is well established. However, the most widely used coarse-grained methods are based on backbone geometry alone and do not take into account the chemical nature of the residues. Elastic Network Contact Model (ENCoM) is a coarse-grained NMA method that includes a pairwise atom-type non-bonded interaction term, which makes it sensitive to the sequence of the studied molecule. We adapted ENCoM to simulate the dynamics of ribonucleic acid (RNA) molecules.

**Results:** ENCoM outperforms the most commonly used coarse-grained model on RNA, Anisotropic Network Model (ANM), in the prediction of b-factors, in the prediction of conformational change as measured by overlap (a measure of effective prediction of structural transitions) and in the prediction of structural variance from NMR ensembles. These benchmarks were derived from the set of all RNA structures available from the Protein Data Bank (PDB) and contain more total cases than previous studies applying NMA to RNA. We thus established ENCoM as an attractive tool for fast and accurate exploration of the conformational space of RNA molecules.

**Availability:** ENCoM is open source software available at <u>https://github.com/NRGlab/ENCoM</u>

## INTRODUCTION

It is now well-established that RNA molecules possess a diverse range of functions in all domains of life and there is growing interest in understanding and characterizing these functions. However, our structural knowledge of RNA is still scarcer than that of proteins, which is based on about two orders of magnitude more structures in the Protein Data Bank (PDB) (1). RNAs are dynamic and thus exist in an ensemble of conformations that are intimately tied to their function (2). The computational prediction of RNA conformational ensembles from a known structure or model is attractive. In the case of proteins, a well-established method for obtaining such information is coarse-grained Normal Mode Analysis (NMA) using Elastic Network Models (ENMs) (3). Many different ENMs with various application niches have been developed for proteins, while the few studies applied to RNA (4, 5) have used the Anisotropic Network Model (ANM) (6).

We recently introduced the Elastic Network Contact Model (ENCoM) for NMA of proteins. It was shown to perform significantly better than other models, including ANM, in the case of conformational change prediction (7). Here, we adapted ENCoM to work on RNA molecules and benchmarked its performance on a set of curated RNA structures derived from the 1194 RNA-only structures available in the PDB resolved by either Nuclear Magnetic Resonance (NMR) or X-ray crystallography. ENCoM outperforms ANM for experimental b-factor prediction, conformational change prediction, and NMR ensemble conformational variance prediction.

Under the approximation that all motions of a macromolecule are harmonic, NMA, whether coarse-grained or all-atom, is an analytical technique that can explore the whole theoretical conformational space of the macromolecule at every timescale with a low computational cost. In contrast, molecular dynamics (MD) simulations make no such approximation, but require massive amounts of computation to explore longer timescales (8). In NMA, the molecule is represented as a system of beads connected by springs with a harmonic potential. For a system of N beads, the normal modes are the 3N eigenvectors of the Hessian matrix, each with its associated eigenvalue. The first six normal modes are trivial motions that represent the translational and rotational degrees of freedom of the centre of mass of the system. From the seventh, the modes are ordered with increasing frequency and thus increasing energetic cost to achieve the same amount of deformation. Usually a small set of the first non-trivial modes is sufficient to describe global motions of biological significance, for example in the case of the hinge-bending motion in citrate synthase (9).

In the study of proteins, the standard coarse-grained ENMs use one mass per amino acid, located on the C_α_ atom (3). There have been efforts to expand these coarse-grained models to include more information, such as the C_β_ atom (10), or the combination of different levels of coarse-graining (11, 12). However, single-mass per residue models continue to be the standard for coarse-grained NMA of proteins and seem to preserve well the nature of the slowest and most global motions. The most widely-used model for proteins is arguably ANM (13), using one mass per residue at the C_α_ atom by default. By contrast, in the two cases where ANM was applied to RNA, the model performed significantly better when three masses were assigned for each nucleotide: one for the phosphate group, one for the sugar group and one for the nucleobase (4, 5). This improvement can be explained in part by the higher intrinsic flexibility of RNA, having six backbone torsional angles per residue instead of three (14), and also by the importance of base pairing in forming specific three-dimensional (3D) architectures. We report similar findings with increased performance of both ENCoM and ANM across all benchmarks when using three masses per residue instead of one.

## MATERIALS AND METHODS

### The ENCoM model

The ENCoM potential is based on a model called generalized Spring Tensor Model (STeM), which uses a potential function containing four terms: covalent bond stretching, angle bending, dihedral angle torsion and non-bonded (or long-range) interactions (7, 15).

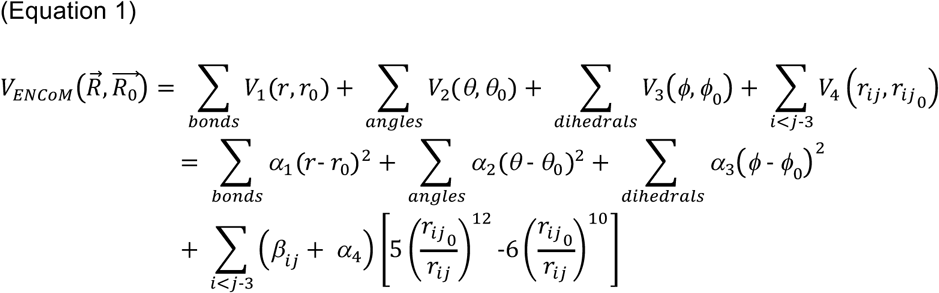

In the ENCoM potential, the non-bonded interaction term is modulated according to the surface area in contact between the residues by the *β*_*ij*_ terms.

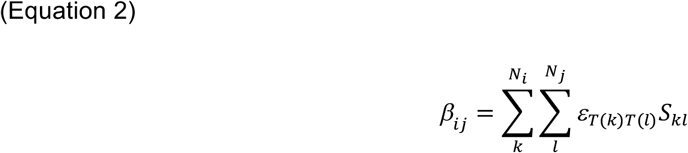

*ε*_*T*(*k*)*T*(*l*)_ represents the interaction between atoms of types T(k) and T(l) while *S*_*kl*_ is the surface area in contact between the two atoms, calculated using a constrained Voronoi procedure, as described by McConkey and co-workers (16). The atom types used are those of Sobolev and co-workers (17), which are divided in eight classes: hydrophilic, acceptor, donor, hydrophobic, aromatic, neutral, neutral-donor and neutral-acceptor. Supplementary Table 1 shows the assigned type of every atom from the four common ribonucleotides.

ANM uses a very simple potential, where each pair of nodes that are closer than a certain distance cutoff are considered to be interacting and are connected with a spring of uniform constant *γ* (18). The interaction distance cutoff *R*_*c*_ (generally set to 18 Angstroms (Å) for proteins) is thus a parameter of the model, which we have extensively tested in increments of 1 Å throughout the different benchmarks. The higher and lower bounds for the cutoffs tested were chosen to show a clear decreasing trend in either direction, confirming that the value for maximal performance has been found.

In the case of proteins, it is generally believed that ENMs using only one node per residue situated on the C_α_ atom still capture the essential low-frequency motions of the molecule (3). However, for RNA molecules that are intrinsically more flexible than proteins, using more nodes per residue leads to an increase in the predictive power of the model (4). We tested both a single-node per residue model (1N model), where the mass is located at the phosphate atom, and a three-nodes per residue model (3N model), where we use one mass each for the sugar, base and phosphate groups positioned on the C1’, C2 and P atoms, respectively, as already used in the case of an ANM (5). Figure 1 shows the positioning of the masses in the four standard nucleotides. The base pairs shown were extracted from a double-helix generated using the MC-Fold and MC-Sym pipeline (19).

**Figure 1.**
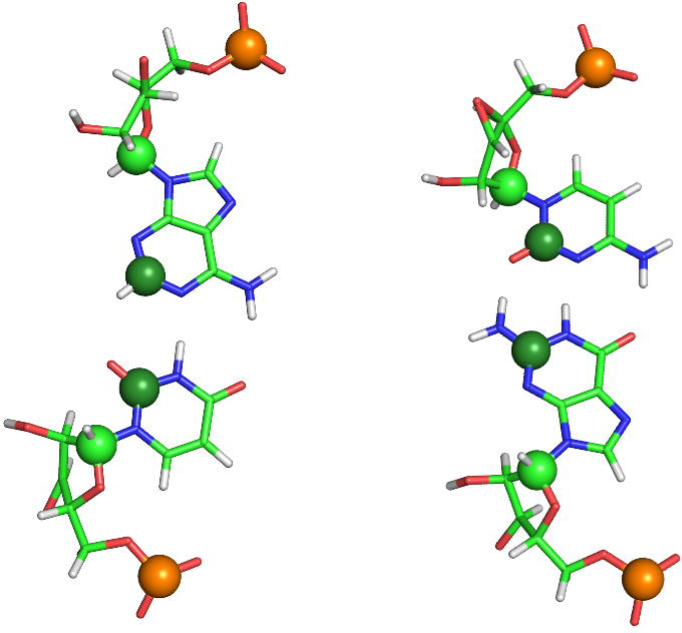
Assignment of the masses on the four standard nucleotides, arranged in two base pairs extracted from a standard double helix. The phosphate atoms are in gold, the C1’ carbons from the sugar group in light green and the C2 carbons from the nucleobase in dark green. All these masses are used in the 3N models while only the phosphate mass is used in the 1N models.

### Dataset of RNA Structures

Experimental determination of RNA structure is achieved by many different techniques, of which two are by far the most common: X-ray crystallography (20) which takes a snapshot of the molecule near its lowest energy conformation and Nuclear Magnetic Resonance (NMR) (21, 22) in which an ensemble of probable conformations is resolved. In the PDB as of 2017-04-20, there were 708 structures solved by X-ray crystallography and 486 NMR ensembles when restricting the search to entries containing only RNA. We used these data in our benchmarks after applying various filters to compare ANM to ENCoM in the prediction of b-factors and conformational transitions. We use all chains present in the biological assembly for the X-ray structures and all chains in the structural ensemble for NMR experiments.

We chose to restrict the model to the four standard nucleobases, replacing all the modified nucleosides with their unmodified analog using the ModeRNA software (23), which contains information about all the 170 modifications present in the MODOMICS database (24). We also added missing atoms where it was the case, for example terminal phosphate groups so that each residue was complete. This addition ensured that the assignment of the masses to the residues was standard for all residues. The structures for which ModeRNA produced an error or which produced an error for any of the models tested subsequently were removed from the analysis. Moreover, the size of the X-ray resolved RNA structures was restricted to a range between 15 and 300 nucleotides inclusively to mitigate potential artefacts of crystal packing in the case of the lower bound, and to reduce computational cost in the case of the upper bound. This left us with 522 X-ray structures in the dataset from the 708 initially present in the PDB. In the case of the NMR resolved RNAs, no size threshold was applied as we trusted the reliability of these experiments even for smaller molecules and no molecules bigger than 155 residues have been elucidated this way. Since we were interested in dynamical ensembles solved by NMR, we restricted our analysis to submissions containing at least two models. This procedure left us with 369 NMR ensembles out of the 486 originally considered, having an average size of 30 nucleotides. The list of all PDB codes of the structures kept for the three benchmarks is given in Supplementary Table 2.

### Clustering

We performed complete linkage clustering (25) on the sequences from our database of structures with a distance threshold of 0.1, ensuring that any pair of sequences within a given cluster were at least 90% similar. The similarity metric used for calculating the distance is derived from the score of a Needleman-Wunsch global alignment (26) between the two sequences, with the following scoring scheme: gap penalty: -1 mismatch penalty: -1 match score: 1.

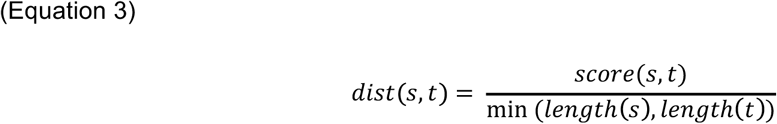

### B-factors

Experimental B-factors can be predicted from calculated normal modes as described before by Atilgan and co-workers (6).

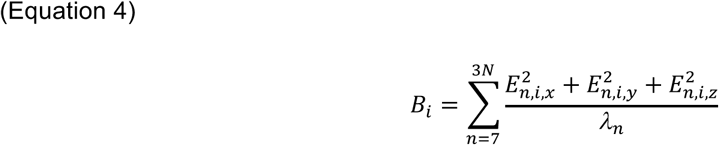

where *E*_*n,i*_ represents the n^th^ eigenvector and *λ* its associated eigenvalue. The Pearson correlation coefficient is then calculated between the predicted and experimental b-factors. In addition to a resolution filter of 2 Å or higher, we used only the structures for which no modified nucleobases were initially present and in which all residues were complete. This restriction was applied to ensure that every atom used in the prediction had a corresponding experimental b-factor, and not an extrapolated value from the rebuilding of the modified nucleobases and missing atoms by ModeRNA. The structures were clustered according to their sequence as described and we computed the mean correlation normalized by cluster.

### Overlap

The overlap between two conformations is a measure of the similarity between the eigenvectors predicted using the start conformation and the displacement vector 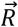 calculated between the coordinates of the target and start conformations, after both were superimposed (9, 27).

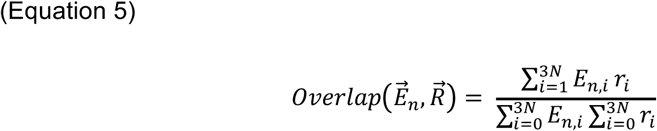

For each eigenvector 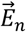, it has a value between 0 and 1, which describes how well we can reproduce the target conformation by deforming the start conformation along the eigenvector. We calculate the best overlap from the first ten eigenvectors, which correspond to the first ten energetically accessible modes.

To ensure that the same conformation is not sampled twice, we reject pairs of conformations for which the root mean squared deviation (RMSD) of the phosphate atoms is lower than 0.75 Å. Clustering was performed as described. We calculated the overlap both ways for every pair from every sequence cluster and took the mean best overlap normalized by cluster. This ensures that molecules which have a lot of conformations sampled in the PDB do not drive the score and instead each RNA or family of RNAs has an equal weight.

### PCA

Principal component analysis (PCA) is a statistical technique that is used to transform observations of variables which may de correlated into linearly uncorrelated variables which are called principal components (PCs) (28). PCA was used here to extract dominant motions apparent within the conformational ensembles obtained from NMR experiments. Each conformation is represented as a vector of length 3N where N is the number of nodes. PCA was computed on these vectors using a singular value decomposition (SVD) algorithm (29). The PCs obtained are analogous to normal modes in that they are the eigenvectors of the covariance matrix of the 3N coordinates of the ensemble of structures. The first PC describes the largest proportion of the variance in the ensemble, and each subsequent component captures the largest proportion of the remaining variance and is orthogonal to the preceding components. The normalized cumulative overlap (NCO) between the first X normal modes and the first Y principal components is given by:

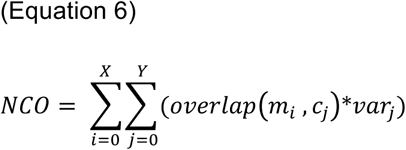

where *m*_*i*_ is normal mode *i, c*_*j*_ is component *j* and *var*_*j*_ is the proportion of variance explained by component *j.* This ensures a value between 0 and 1 as both the normal modes and the components are orthogonal, and *var_j_* sums to 1 over all the components. For the current study, we used the three first non-trivial normal modes and the first ten PCs. The normal modes were computed from the conformation of minimal energy in each NMR ensemble, which by convention is the first one in the experimenters’ submission. Clustering was performed as described and we computed the mean NCO normalized by cluster.

### Bootstrapping

Bootstrapping is a statistical resampling technique that allows the estimation of the variance of a metric by resampling observations from the original sample with replacement, over many iterations (30). in our benchmarks, we used the bootstrapped mean values over 10,000 iterations as the final performance value of the models, with the standard deviation from the bootstrap as the estimated error. This technique compensates for the small sized datasets and gives a better basis for comparing the different models.

### Analysis of RNA families

To see if the models perform better on certain types of RNAs, the sequence of every RNA structure used in the benchmarks was submitted to the Rfam web Application Programming Interface (API) (31). No sequence matched more than one family, and the ones that matched none were not considered for this analysis. For all three benchmark, the mean performance of each model was given for every Rfam identified.

## RESULTS

### Experimental b-factor prediction

X-ray crystallography gives rise to experimental temperature factors commonly called b-factors, which measure how much each atom oscillates around its equilibrium position in the crystal. The correlation between these fluctuations and those predicted by the ENMs is a common benchmark, which we have performed here on high-resolution X-ray RNA structures of 2 Å or less. The clustering step left us with 18 high-resolution structures divided in 12 sequence clusters. Figure 2 gives the Pearson correlation between the predicted and experimental B-factors. As expected, the optimal cutoff for ANM is higher for the 1N than for the 3N versions at respectively 9 and 29 Å, as individual nodes are on average further apart when there is only one node per residue.

**Figure 2.**
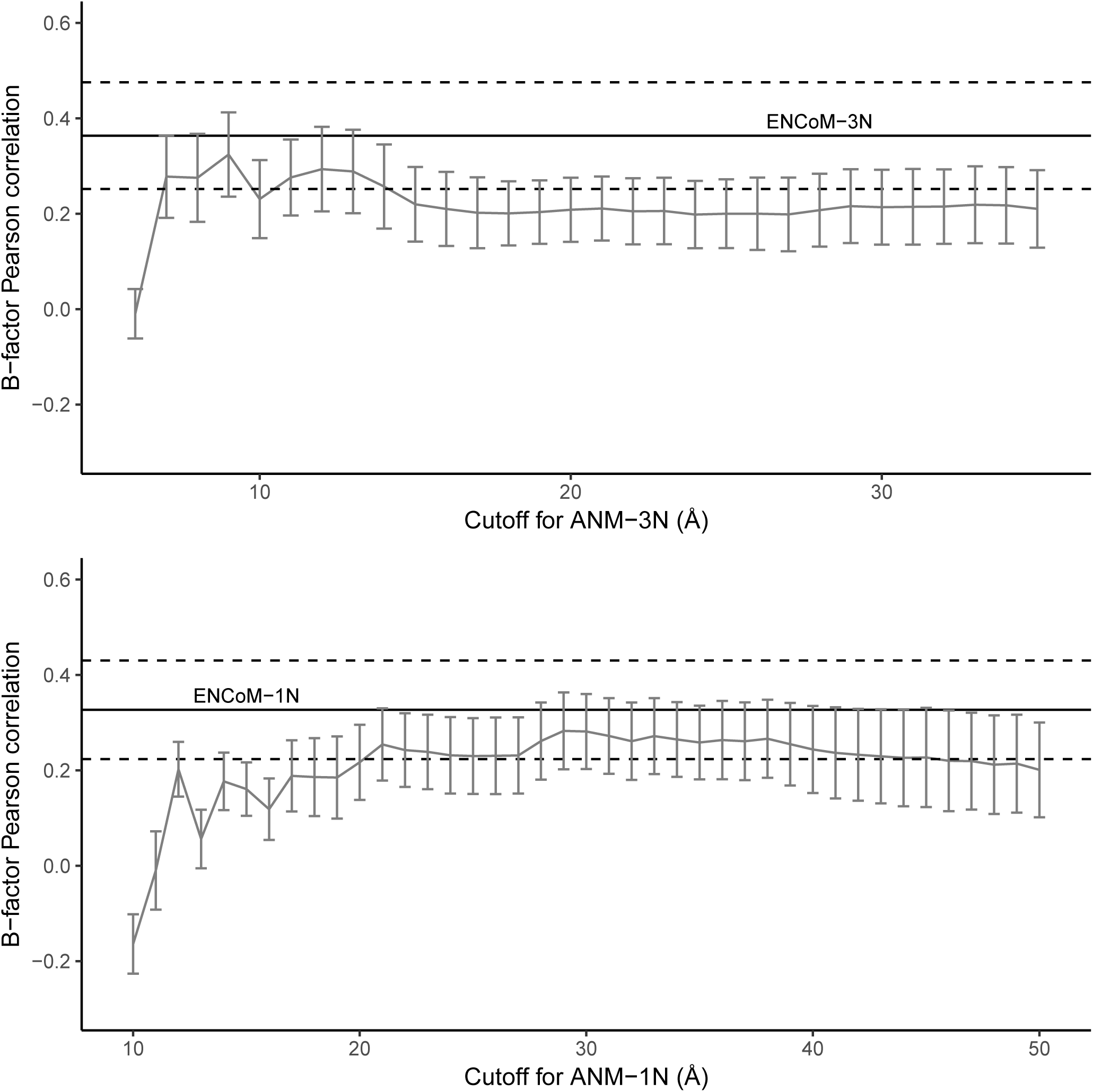
Pearson correlation coefficient between predicted and experimental B-factors for ENCoM and ANM. 18 high resolution (2 Å or less) RNA crystal structures were used and the results were bootstrapped over 10,000 iterations. Similar sequences were clustered before bootstrapping to prevent over-representation of a single molecule, giving 12 clusters at the threshold of 90% sequence identity. The mean of each cluster was used in the resampling and the estimated standard deviation is shown with error bars in the case of ANM; dashed lines in the case of ENCoM. Results are shown for both 1N and 3N models. The interaction distance cutoff for ANM was varied in steps of 1 Å. ENCoM does not have any user-defined parameters, except for the number of nodes per residue, and so a horizontal line summarizes the results. ENCoM performs better than ANM at every interaction distance cutoff tested.

The highest correlation among the models tested is 0.36 for ENCoM-3N, which is low when compared to the correlations previously obtained with ENCoM and other tools applied to proteins (7), all between 0.54 and 0.60. In addition, b-factor correlations of up to around 0.6 have been reported with ANM applied on RNA in a similar fashion as here, and with 3 masses per residue by Zimmerman and Jernigan (4). However, we could not verify their results as their paper does not mention what structures they used and the website containing the supplementary data (including the structures) is not operational. The mean correlation from each of the 12 sequence clusters for both ENCoM-3N and the best ANM-3N is given in Supplementary Table 3. Interestingly, for three individual sequence clusters the correlations obtained for ENCoM-3N and ANM-3N models are negative.

### Conformational change prediction

Another source of information regarding dynamics arising from X-ray crystallography is the crystallization of the same molecule in different conformations due to either varying experimental conditions or the presence of slight mutations in the sequence of residues. After clustering and applying filters (see Materials and Methods), we obtain 185 structures divided in 32 sequence clusters. Each of these clusters contains at least one pair of distinct conformations for a given RNA. We use these pairs to benchmark the ability of the models to predict conformational change. This ability is in many ways more relevant to a biological context, as the conformational variation resulting from different experimental conditions happens on longer timescales than b-factors and can have functional significance (8).

Figure 3 presents the average best overlap from pairs of different conformations, computed from the ten first modes. This metric gives a general idea of how well these first modes describe the conformational change, but a value near 1 was not expected since this is not an additive measure (Eq. 5). Here, the 3N versions for both ENCoM and ANM have a significant advantage over the 1N versions. Again, for ANM the optimal interaction distance cutoff is higher for the 1N than for the 3N version. The best performance is again obtained by ENCoM-3N at 0.47 mean best overlap, followed by ANM-3N at 9 Å cutoff with 0.44 overlap, a whole bootstrap standard deviation lower as can be clearly seen on the graph (Fig. 3). This result corresponds to our previous results benchmarking ENCoM on proteins, where ENCoM performed highest of all the models tested on the conformational change benchmark (7).

**Figure 3.**
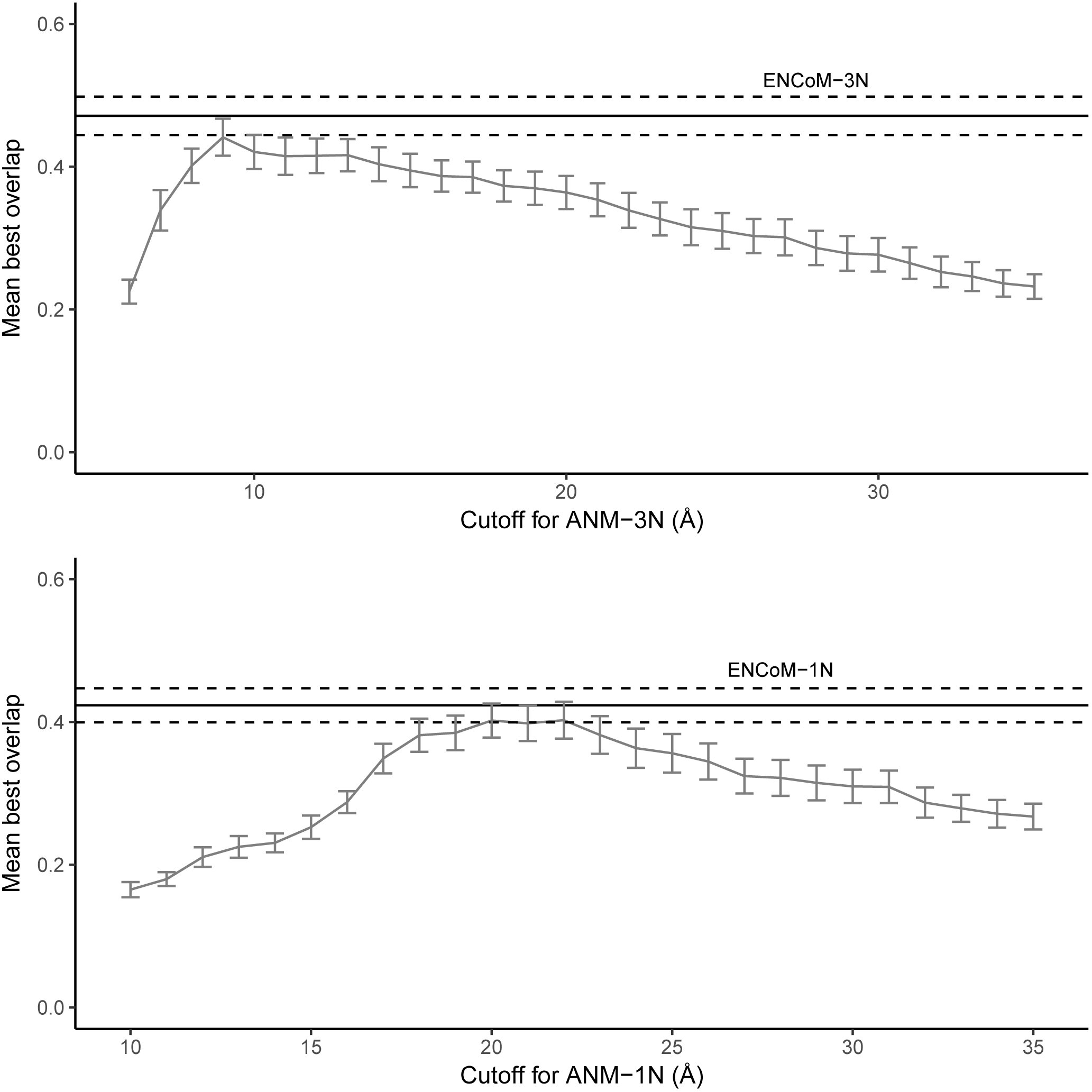
Overlap between predicted and observed conformational changes. Average best overlap between two different conformations of the same molecule calculated both ways from the first 10 modes. The same molecule is defined as 90% sequence identity, and only the paired residues in the alignment are used for computing the overlap. 185 crystal structures were compared in duos both ways, with 32 clusters of minimal intra-cluster sequence identity of 90%. Results were bootstrapped over 10,000 iterations.

Figure 4 shows in greater detail the distribution of average best overlap for the two best performing models: ENCoM-3N and ANM-3N with 9 Å cutoff. The resampling was also done over 10,000 iterations, but at each iteration a random pair of conformations from every cluster was selected. The mean from these resamples was computed at every iteration, and these 10,000 resampled means are shown as histograms (Fig. 4). We can reject the null hypothesis that ENCoM has no advantage over ANM on the prediction of conformational changes with a p-value of 0.027.

**Figure 4.**
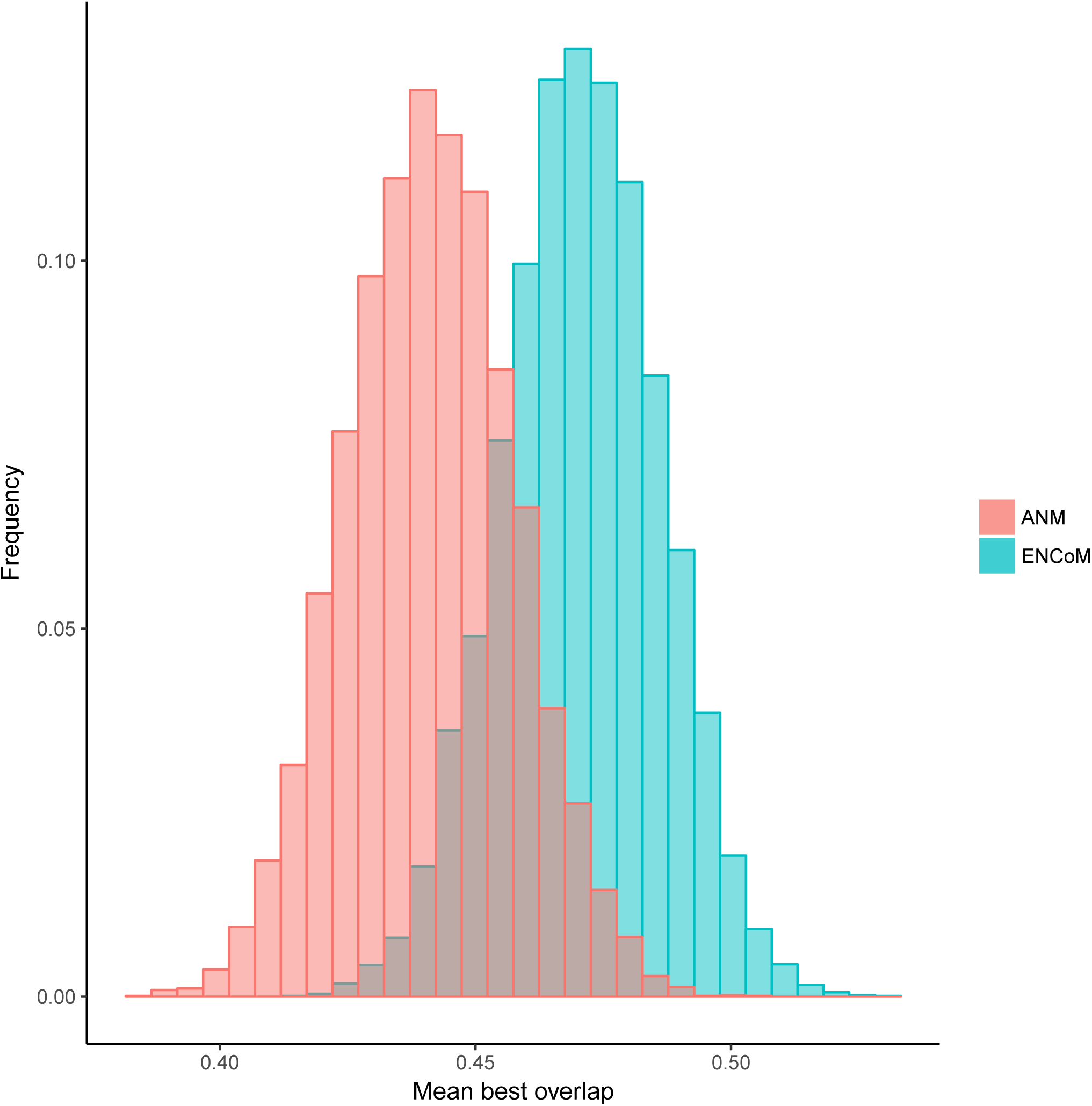
Comparison of mean best overlap between ENCoM and ANM using 3 nodes per residue. The best ANM interaction cutoff among those tested (9 Å) was used. The same 185 crystal structures grouped in 32 clusters were used. 10,000 iterations of bootstrap were performed, selecting one pair from each cluster at each iteration. The 10,000 mean best overlaps obtained are shown as a histogram. The null hypothesis that the mean difference between the two Gaussians is 0 is rejected with p = 0.027.

### Overlap with NMR ensemble variance

In contrast to X-ray crystallography, NMR experiments give us direct information about the conformational space of the molecule studied over vast timescales. The motions apparent within the structural ensembles resolved using this technique can inform us on the ability of NMA models to recreate the conformational space of a molecule from its lowest energy conformation. After clustering as described, we obtained 369 NMR ensembles separated in 288 sequence clusters.

Figure 5 shows the NCO of the first 10 slowest modes with the principal component analysis motions derived from NMR conformational ensembles (Eq. 6). Here again, the optimal cutoff for ANM- 3N is lower than the one of ANM-1N at 10 Å versus 20 Å. The best performing model is ENCoM-3N with a mean cumulative overlap of 0.40 and the second-best is ANM-3N with 10 Å cutoff at 0.37. These relatively low values could be explained by the small size of molecules solved by NMR compared to molecules solved by X-ray, and the fact that coarse-grained NMA models tend to perform better on bigger molecules because they are best at describing cooperative movements.

**Figure 5.**
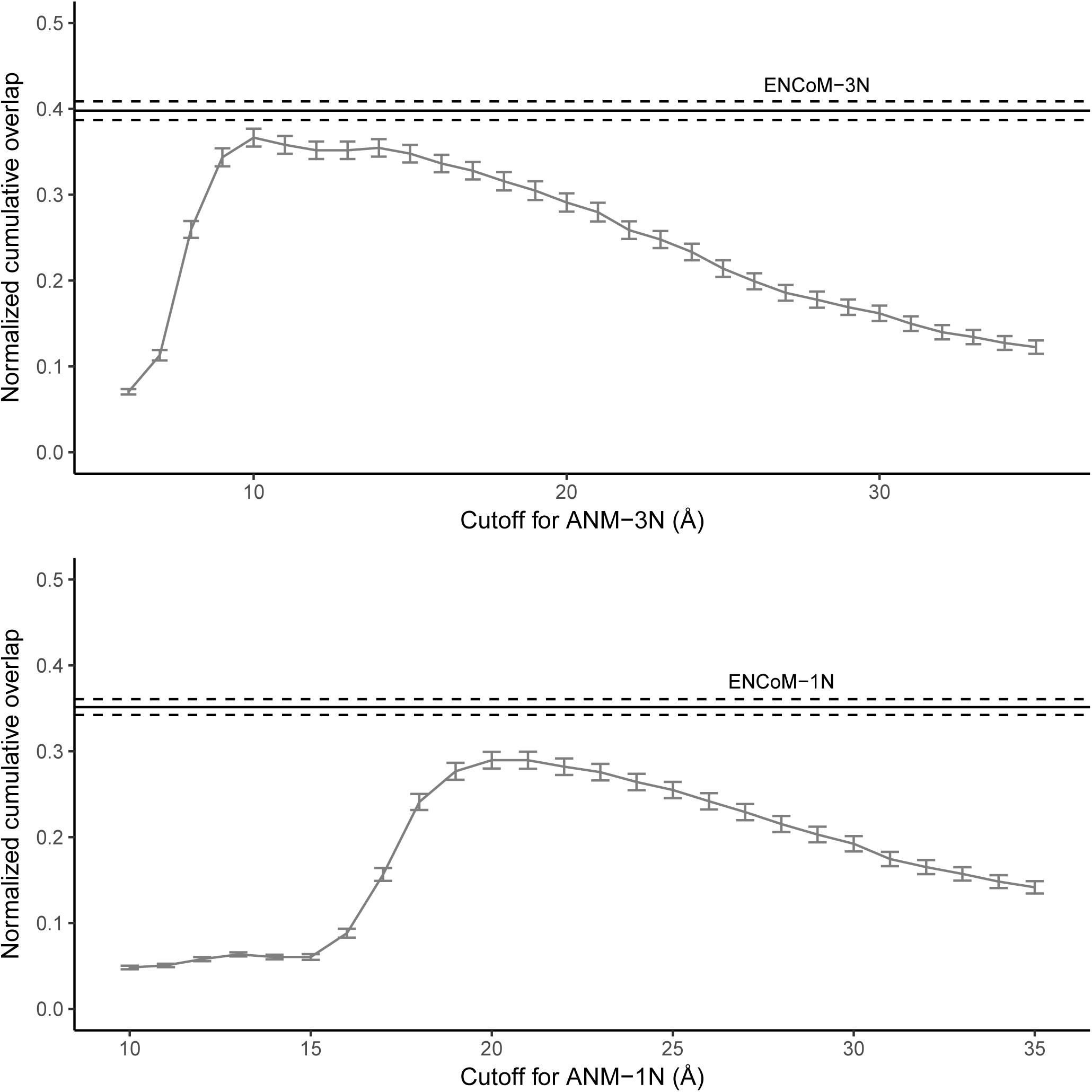
Normalized cumulative overlap (NCO) of the first 10 modes predicted from the least energy structure of NMR ensembles and the 10 first principal components (PCs) generated from principal component analysis on the whole ensemble. 369 structural ensembles were analysed and grouped in 288 clusters of 90% or more intra-cluster sequence identity. Results were bootstrapped over 10,000 iterations.

### Rfam pooling

For the three benchmarks outlined above, we pooled the sequences that matched specific RNA families together and showed the performances of ENCoM and ANM in both 1N and 3N versions. Table 1 reports the results for b-factor prediction; Table 2 for conformational change prediction; and, Table 3 for NCO of PCA from NMR ensembles. In each table, the cutoffs used for the two ANM models were those giving the highest performance for the particular benchmark reported.

In Table 1, only three Rfams were identified due to the small size of our initial dataset. For these three families, the best model overall is ANM-3N with 0.53 b-factor Pearson correlation, followed by ENCoM-3N at 0.50. These results are both significantly better than the average performance reported in Figure 2, which indicates that the three aberrant cases with negative correlations previously outlined do not correspond to annotated Rfams. It is important to note that the sample size is small here, and that although ANM-3N performs better overall, ENCoM obtains the two highest mean correlations among all families, in the case of the purine riboswitch family.

For the conformational change prediction benchmark summarized in Table 2, it is interesting to note that while ENCoM-3N performs better overall, ANM is significantly better for some individual families. Some of these families are single cases, but others are quite populated like the tRNA family with 8 cases. The purine riboswitch with 26 cases is also interesting as the performances of ENCoM- 3N and ANM-1N are tied.

Table 3 shows the NCO of PCA from NMR ensembles. While results for individual families vary, the average performance seems to indicate that ENCoM has a slight advantage over ANM at 0.31 for ENCoM-3N compared to 0.29 for ANM-3N. This result is lower than the 0.40 average reported in Figure 4 and is driven down by two families for which all models perform poorly: Gag/pol translational readthrough site and U6 spliceosomal RNA. Both of these families contain a single case and the poor performance may be specific to the experiments conducted.

## DISCUSSION

### Overall performance

We have shown here the superior performance of ENCoM over ANM when applied to RNA across three extensive benchmarks, namely experimental and predicted B-factors correlation, conformational change prediction, and NMR ensemble variance prediction. When looking at average performance over all structures tested a clear trend emerges that using three masses per residue placed as Pinamonti and co-workers (5) gives a performance boost to both models. Among these 3N models, ENCoM performs better on average by a significant margin.

### Artifacts from crystallographic b-factors

For three of the twelve sequence clusters used in the experimental b-factor prediction benchmark, both ENCoM and ANM get negative correlations (Table S3). This means that both models predict regions that are rigid according to the experiments as flexible, and vice versa. If we exclude the cases from these three clusters as experimental flaws and take the mean of the 9 remaining clusters, we get a correlation of 0.56 for ENCoM-3N and of 0.46 for the best ANM-3N, which are much closer to what is observed when applying NMA models to proteins (7). It would be interesting to investigate in more details the nature of the anti-correlations to see if some factor in the experimental conditions or the nature of the RNAs could explain the discrepancy.

However, in our case, we are more interested in predicting conformational changes than b-factors, and we have previously shown that parameter optimization offers a trade-off between conformational space and b-factors prediction (7). Consequently, we did not investigate further this observation. Furthermore, b-factors have been shown to adopt biphasic behavior with a different slope of variation for cryogenic versus standard temperatures (32). Since most X-ray crystallography experiments are conducted at cryogenic temperatures, their b-factor measurements are thus less relevant to biological function. B-factors can arise from crystal packing and rigid body motions which are also irrelevant in a biological context (33).

### Parameter-free conformational change prediction

The best performance in the conformational change prediction benchmark was obtained by ENCoM- 3N with 0.47 average best overlap. This result falls in between the values previously obtained (7) for domain and loop motions of proteins, respectively around 0.62 and 0.35. Such a performance indicates that the motions observed in our database of RNA conformational changes are not as concerted as domain motions, but more so than loop motions on average. It is important to note that the advantage over ANM reported here is slighter than previously reported on proteins. However, it is most probably a result of exploring the interaction cutoff extensively and comparing to the best performing value. In a real case scenario, if one were to use ANM, it would be very hard to properly estimate the best cutoff to use to get the most realistic prediction power for conformational changes. Even if ANM performed on par with ENCoM, ENCoM would still have the advantage of being parameter-free.

### tRNA family

When looking at individual RNA families, we encounter some exceptions in terms of the best performing models, as outlined in Table 1. ANM performs notably better for the tRNA family which contains eight individual structures in our database for the conformational change prediction benchmark. The best performing model for this family is ANM-1N at 0.45 mean best overlap and the second-best is ENCoM-1N with 0.44 mean best overlap. In addition to ANM performing better, it is important to note that for this family no advantage seems to be conferred by the usage of three nodes per residue. It is widely known that tRNA molecules contain a wide variety of modified nucleobases (34). These bases confer specific properties to the anticodon, but are also thought to play structural roles in the folding of tRNAs. Since we explicitly replace the modified bases with their standard analog in the present study, we likely introduce noise that negatively affects ENCoM as the surface complementarity measured in the interactions where a modified base is implicated is different than the real surface. However the ANM potential depends only on the geometry of the initial structure and such details do not change its predictions.

### Purine riboswitches

The purine riboswitch family in the conformational change benchmark contains 26 members and the conformational changes encompassed are best represented by both ENCoM-3N and ANM-1N at 22 Å interaction distance cutoff, with 0.48 mean best overlap. It is possible that the absence of metal ions in our models, which have been shown to have important effects in determining the conformation of these riboswitches (35), is negatively impacting the performance of ENCoM while leaving ANM unaffected. Since ENCoM’s advantages rely on the atomic surface complementarity term, the absence of these ions means that local interactions between the negatively charged backbone and the positive ions are not accounted for, and the advantage is lost. As before, ANM would not suffer from such inaccuracies, as the result is completely determined by the geometry of the starting conformation.

### Parameterization

The parameterization of ENCoM was done for proteins and while it was shown to be quite robust, even to changes in the interaction matrix of the different atom types (7), we were not expecting it to perform initially well on RNA. However, since this was the case, we did not delve into parameter optimization for the present study, but we would be interested in doing so in the future. In contrast, ANM depends on the interaction distance cutoff, for which the value giving best performance varies according to the benchmark. This seems to indicate that for blind prediction ENCoM would be the preferred model.

## Conclusions

We presented a version of ENCoM specifically tailored to work optimally on RNA molecules by allowing the use of three masses per nucleotide in the coarse-graining of the input structure. We have shown that it consistently outperforms ANM across all benchmarks in addition to being parameter-free as far as the user is concerned. Another advantage of ENCoM is the possibility to estimate the effect of mutations, which ANM is incapable of due to its sequence-agnostic nature. However, ENCoM is more sensitive to inaccuracies in the input structures such as the rebuilding of modified bases or the presence of metallic ions, which are not currently taken into account.

## AVAILABILITY

ENCoM is open source software available in the GitHub repository (<u>https://github.com/NRGlab/ENCoM</u>)

## SUPPLEMENTARY DATA

Supplementary Data are available at NAR online.

## ACKNOWLEDGEMENTS

We thank Marie Papineau, Frédéric Mailhot and Etienne Richan for their careful reading of the manuscript. RJN is part of PROTEO (the Québec network for research on protein function, structure and engineering) and GRASP (Groupe de Recherche Axé sur la Structure des Protéines).

## TABLES AND FIGURES LEGENDS

**Table 1. Performance comparison for b-factor Pearson correlation for ENCoM and ANM for different Rfam RNA families**. The mean performance measure for the Rfam is given for both the 1N and 3N versions of the models. The best interaction cutoffs for ANM-1NN are used, respectively 29 Å and 9 Å. The best performing model for each Rfam is shown in bold and the average over all Rfams is shown on the last line.

**Table 2. Performance comparison for mean overlap between the best of the 10 first modes and an experimentally observed conformational change, for ENCoM and ANM for different Rfam RNA families**. The mean performance measure for the Rfam is given for both the 1N and 3N versions of the models. The best interaction cutoffs for ANM-1N and ANM-3N are used, respectively 22 Å and 9 Å. The best performing model for each Rfam is shown in bold and the average over all Rfams is shown on the last line.

**Table 3. Performance comparison for normalized cumulative overlap of the first 10 modes and the first 10 PCs from PCA on NMR ensembles, for ENCoM and ANM for different Rfam RNA families**. The mean performance measure for the Rfam is given for both the 1N and 3N versions of the models. The best interaction cutoffs for ANM-1N and ANM-3N are used, respectively 20 Å and 10 Å. The best performing model for each Rfam is shown in bold and the average over all Rfams is shown on the last line.

## SUPPLEMENTARY DATA LEGENDS

**Supplementary Table 1. Atom type assignation of the four standard nuclotides.** The atom types assigned are the ones from Sobolev and coworkers, divided in eight classes: hydrophilic, acceptor, donor, hydrophobic, aromatic, neutral, neutral-donor and neutral-acceptor.

**Supplementary Table 2. The PDB codes of the structures used for the three benchmarks.** 18 structures were retained for the prediction of experimental b-factors, 185 for the prediction of conformational changes and 369 structural ensembles for the prediction of NMR ensemble variance.

**Supplementary Table 3. Pearson correlation between predicted and experimental b-factors for each of the 12 sequence clusters.** The mean of every cluster is shown for ENCoM-3N and ANM-3N 9 Å cutoff (the best performing ANM).

